# Genome-wide association meta-analysis identifies novel *GP2* gene risk variants for pancreatic cancer in the Japanese population

**DOI:** 10.1101/498659

**Authors:** Yingsong Lin, Masahiro Nakatochi, Hidemi Ito, Yoichiro Kamatani, Akihito Inoko, Hiromi Sakamoto, Fumie Kinoshita, Yumiko Kobayashi, Hiroshi Ishii, Masato Ozaka, Takashi Sasaki, Masato Matsuyama, Naoki Sasahira, Manabu Morimoto, Satoshi Kobayashi, Taito Fukushima, Makoto Ueno, Shinichi Ohkawa, Naoto Egawa, Sawako Kuruma, Mitsuru Mori, Haruhisa Nakao, Yasushi Adachi, Masumi Okuda, Takako Osaki, Shigeru Kamiya, Chaochen Wang, Kazuo Hara, Yasuhiro Shimizu, Tatsuo Miyamoto, Yuko Hayashi, Yasuyuki Hosono, Hiromichi Ebi, Tomohiro Kohmoto, Issei Imoto, Yoshinori Murakami, Masato Akiyama, Kazuyoshi Ishigaki, Koichi Matsuda, Makoto Hirata, Katsuaki Shimada, Takuji Okusaka, Takahisa Kawaguchi, Meiko Takahashi, Yoshiyuki Watanabe, Kiyonori Kuriki, Aya Kadota, Kenji Wakai, Taiki Yamaji, Motoki Iwasaki, Norie Sawada, Shoichiro Tsugane, Kengo Kinoshita, Nobuo Fuse, Fumiki Katsuoka, Atsushi Shimizu, Satoshi S. Nishizuka, Kozo Tanno, Ken Suzuki, Yukinori Okada, Momoko Horikoshi, Toshimasa Yamauchi, Takashi Kadowaki, Teruhiko Yoshida, Fumihiko Matsuda, Michiaki Kubo, Shogo Kikuchi, Keitaro Matsuo

## Abstract

The etiology of pancreatic cancer remains largely unknown. Here, we report the results of a meta-analysis of three genome-wide association studies (GWASs) comprising 2,039 pancreatic cancer cases and 32,592 controls, the largest sample size in the Japanese population. We identified 3 (13q12.2, 13q22.1, and 16p12.3) genome-wide significant loci (*P*<5.0×10^-8^) and 4 suggestive loci (*P*<1.0×10^-6^) for pancreatic cancer. Of these risk loci, 16p12.3 is novel; the lead SNP maps to rs78193826 (odds ratio (OR)=1.46, 95% CI=1.29-1.66, *P*=4.28×10^-9^), an Asian-specific, nonsynonymous *glycoprotein 2* (*GP2*) gene variant predicted to be highly deleterious. Additionally, the gene-based GWAS identified a novel gene, *KRT8*, which is linked to exocrine pancreatic and liver diseases. The identified *GP2* gene variants were pleiotropic for multiple traits, including type 2 diabetes, hemoglobin A1c (HbA1c) levels, and pancreatic cancer. Mendelian randomization analyses corroborated causality between HbA1c and pancreatic cancer. These findings suggest that *GP2* gene variants are associated with pancreatic cancer susceptibility in the Japanese population, prompting further functional characterization of this locus.

## Introduction

With approximately 33,000 related deaths every year, pancreatic cancer is the fourth leading cause of cancer deaths in Japan, after lung, colorectal, and stomach cancers^1^. The incidence and mortality rates of pancreatic cancer have increased steadily over the past decades, while those of other gastrointestinal cancers have shown a decreasing trend^1^. Despite the increasing burden levied by pancreatic cancer, few modifiable risk factors other than smoking and type 2 diabetes mellitus (T2D) have been identified, and the 5-year survival rates remain the worst (<10%) among major malignancies.

Genome-wide association studies (GWASs) have increasingly revealed the role of inherited genetic variations in pancreatic cancer susceptibility. Since the first GWAS, conducted by the PanScan consortium, identified common variants in the gene coding for the ABO blood group system in 2009,^2^ at least 23 genome-wide significant susceptibility loci have been linked to pancreatic cancer risk^3^. However, fewer loci have been identified for pancreatic cancer than for other common cancers, including breast and colorectal cancers^4,5^. Furthermore, the identified risk variants explained approximately 13% of the total heritability on the basis of GWAS-identified SNPs in a population of European ancestry^6^. These observations suggest that additional risk loci can be identified by increasing the sample size, as evidenced by the trend in the numbers of novel variants reported by PanScan. It is also important to expand the GWAS to populations of non-European ancestry because of differences in minor allele frequencies (MAFs) and patterns of linkage disequilibrium (LD) across diverse populations^7^. In fact, previous GWASs focusing exclusively on populations of Eastern Asian ancestry led to the identification of new susceptibility loci for breast and colorectal cancers^8, 9^.

The majority of the risk loci for pancreatic cancer were discovered in the PanScan GWASs, which included populations of European ancestry. Only two GWASs have been conducted in East Asian populations: one in China^10^ and one in Japan^11^. A total of 8 risk loci (5 genome-wide significant loci and 3 loci with suggestive evidence of association) have been identified for pancreatic cancer, but these loci were not replicated in a previous study using samples from European populations^12^. Therefore, the role of common susceptibility loci in East Asian populations remains uncertain and needs further exploration. To detect additional susceptibility loci for pancreatic cancer, we conducted another GWAS in the Japanese population and then performed a meta-analysis combining all published and unpublished GWAS data in Japan.

## Results

After imputation and quality control, we performed a meta-analysis of three Japanese GWASs comprising 2,039 cases and 32,592 controls as well as 7,914,378 SNPs (**Supplementary Table 1 and Table 2**). Genomic control adjustment was not applied because there was little evidence of genomic inflation (lambda=1.03, **Supplementary Figure 1**). We observed genome-wide significant (*P*<5.0×10^-8^) association signals at 3 loci (13q12.2, 13q22.1, and 16p12.3) (**Figure 1 and Table 1**), for which the genes nearest the lead SNP were *PLUT (PDX1-AS1), KLF5*, and *GP2*, respectively. In addition, 4 loci (1p13.2 (*WNT2B*), 2p12 (*CTNNA2*), 3p12.3 (*ROBO2*), and 9q34.2 (*ABO)*) showed suggestive evidence of association (*P*<1.0×10^-6^) (**Figure 1 and Table 1**). The association results of each study are shown in **Supplementary Table 3.** Among these risk loci, 16p12.3 is a novel locus, with genome-wide significant associations observed for 10 SNPs in this region (rs78193826, rs117267808, rs73541251, rs4609857, rs4544248, rs4632135, rs4420538, rs73541271, rs4383154, and rs4383153) (**Supplementary Table 4**). The ORs for these variants ranged from 1.43 to 1.47, indicating stronger associations in this region than the associations indicated by the odds ratios (ORs) for variants identified in previously published GWASs. The lead SNP maps to rs78193826, a nonsynonymous variant of the *GP2* (**Figure 2**) gene. The risk increased by 46% per copy of the minor T allele (OR=1.46, 95% CI=1.29-1.66, *P*=4.28×10^-9^) (**Table 1**). Regional association plots for the other loci are shown in Supplementary Figure 2. According to the 1000 Genomes Project Phase 3 database, rs78193826 is polymorphic, with a MAF ranging from 3.9% to 6.6% in Asian populations, compared with the much lower MAF (<0.1%) in other human populations (**Supplementary Table 4**). LD maps of these 10 SNPs at 16p12.3 are shown in **Supplementary Figure 3.** Complete LD between 9 of these SNPs (all except rs4420538) was observed in the Japanese population. Among the 10 SNPs in this region, only rs4383153 had available association summary statistics in the previous PanScan publications^2,13^, but this SNP was not significantly associated with pancreatic cancer risk (**Supplementary Table 5**). The functional annotation results for the 10 SNPs at 16p12.3 are shown in **Supplementary Table 4.** The lead SNP rs78193826 was classified as “damaging” according to the Sifting Intolerant from Tolerant (SIFT) algorithm and as “possibly damaging” by Polymorphism Phenotyping v2 (PolyPhen-2). Moreover, the estimated combined annotation-dependent depletion (CADD) score was 20.3. For replication, we selected 4 SNPs (rs78193826, rs73541251, rs117267808, rs4632135) that met either of the following criteria: 1) exonic SNP or 2) intronic SNP with a score of 3 or less according to the Regulome DB database.

**Figure 1.**
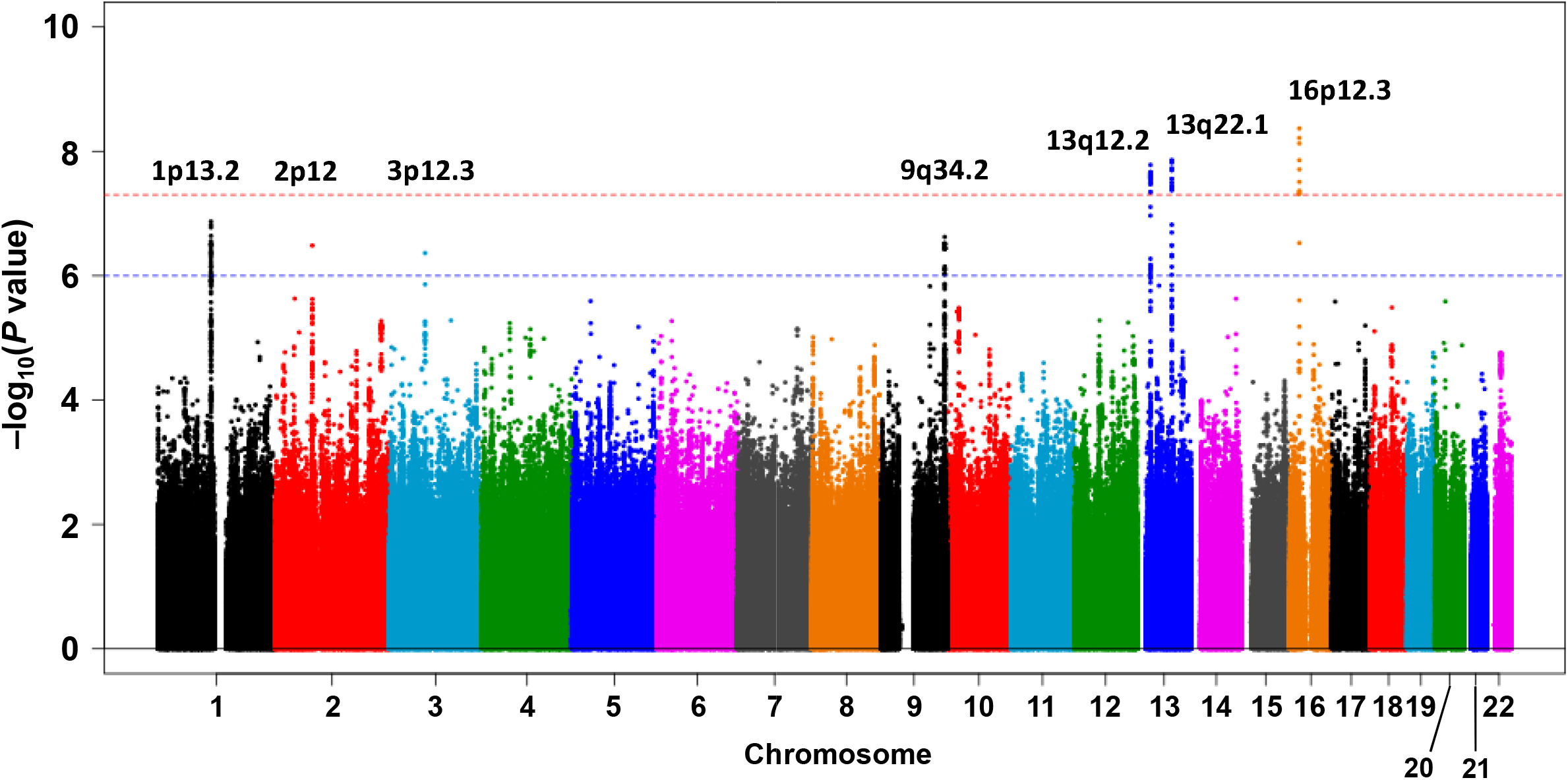
Manhattan plot for the meta-analysis. The horizontal red line represents the genome-wide significance level (α = 5 × 10^−8^). The horizontal blue line represents the suggestive significance level (α = 1 × 10^−6^).

**Figure 2.**
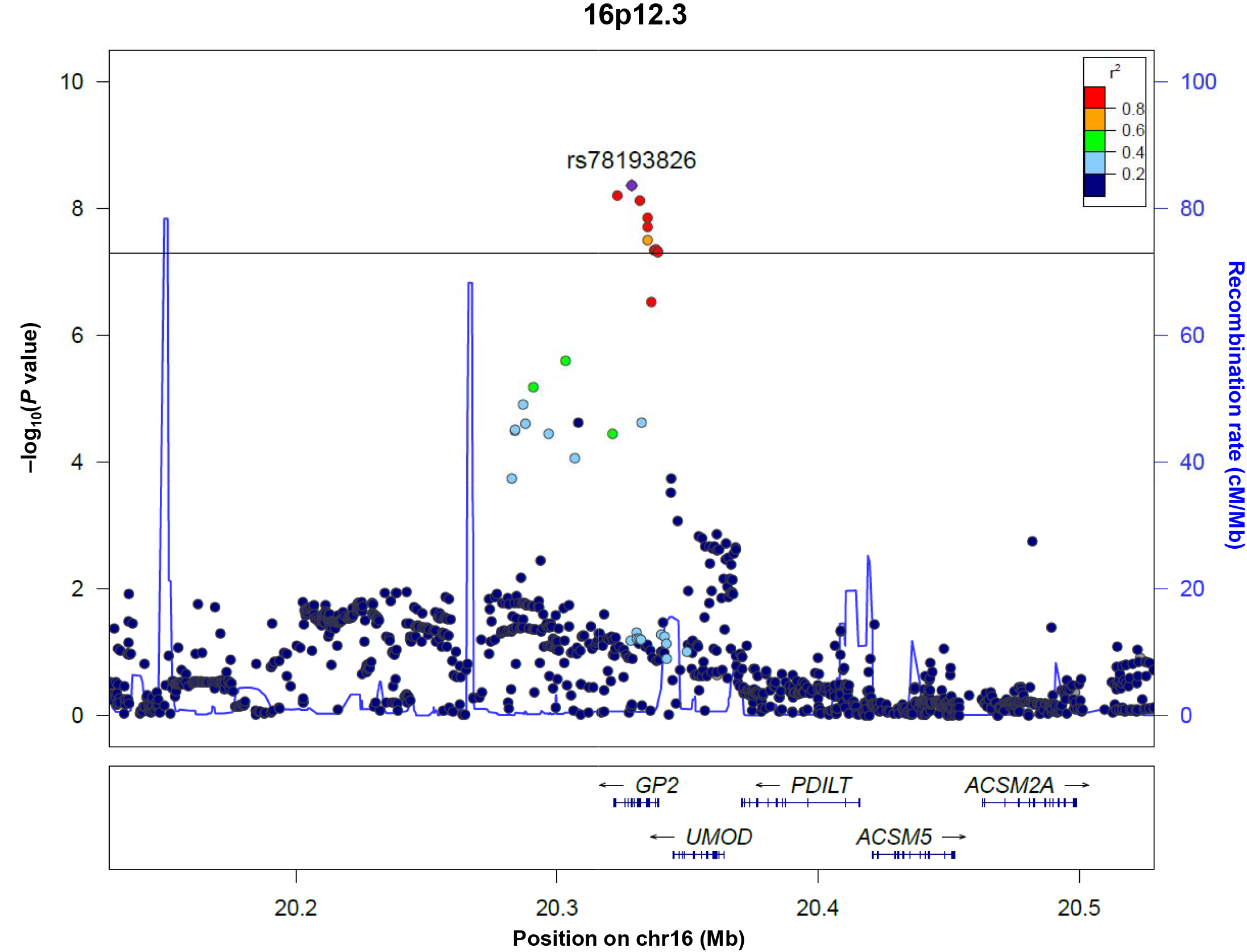
Regional association plot for the 16p12.3 locus identified in the meta-analysis. The vertical axis indicates the −log_10_(*P* value) for the assessment of the association of each SNP with pancreatic cancer. The colors indicate the LD (*r*^2^) between each sentinel SNP and neighboring SNPs based on the JPT population in the 1000 Genomes Project Phase 3.

**Table 1.** Genomic region and lead SNPs associated with pancreatic cancer susceptibility in the meta-analysis of three Japanese GWASs

| SNP | Locus | Chr | Position | Gene | Alleles |  | OR (95% CI) | P value | *I*<sup>2</sup> | HetP value |
| --- | --- | --- | --- | --- | --- | --- | --- | --- | --- | --- |
|  |  |  |  |  | Risk | Non-risk |  |  |  |  |
| Genome-wide significant loci |  |  |  |  |  |  |  |  |  |  |
| rs147905965 | 13q12.2 | 13 | 28474234 | *PLUT* | AT | A | 1.29 (1.18-1.41) | 1.66×10<sup>-8</sup> | 28.3 | 0.248 |
| rs9543325 | 13q22.1 | 13 | 73916628 | *KLF5, LINC00392* | C | T | 1.24 (1.15-1.33) | 1.38×10<sup>-8</sup> | 24.9 | 0.264 |
| rs78193826 | 16p12.3 | 16 | 20328666 | *GP2* | T | C | 1.46 (1.29-1.66) | 4.28×10<sup>-9</sup> | 14.5 | 0.310 |
| Suggestive loci |  |  |  |  |  |  |  |  |  |  |
| rs3737136 | 1p13.2 | 1 | 113060432 | *WNT2B* | G | A | 1.24 (1.14-1.34) | 1.35×10<sup>-7</sup> | 74.3 | 0.020 |
| rs35067842 | 2p12 | 2 | 79568318 | *REG3A, LOC101927987* | C | A | 1.24 (1.14-1.34) | 3.28×10<sup>-7</sup> | 0 | 0.883 |
| rs6809193 | 3p12.3 | 3 | 76479914 | *ZNF717, ROBO2* | G | A | 1.23 (1.14-1.33) | 4.33×10<sup>-7</sup> | 34.2 | 0.219 |
| rs7855466 | 9q34.2 | 9 | 136121303 | *OBP2B, ABO* | T | C | 1.22 (1.13-1.31) | 2.38×10<sup>-7</sup> | 56.7 | 0.100 |

OR values represent the increased risk of pancreatic cancer per risk allele copy for each SNP. Chr, chromosome. RAF, risk allele frequency.

Analysis of another independent replication cohort comprising 507 cases and 879 controls showed that rs4632135 (an intronic variant) was nominally significantly associated with pancreatic cancer risk (*P*<0.05), whereas the other 3 SNPs did not show nominal significance (**Table 2**). However, the direction and magnitude of the effects for all 4 SNPs was consistent with those observed in the GWAS meta-analysis. Furthermore, the combined analysis of the three GWASs and the replication dataset yielded lower *P* values than those of the GWAS meta-analysis for each SNP (**Table 2**), suggesting that the novel risk locus at 16p12.3 discovered in our GWAS meta-analysis was unlikely to be false positive.

**Table 2.**
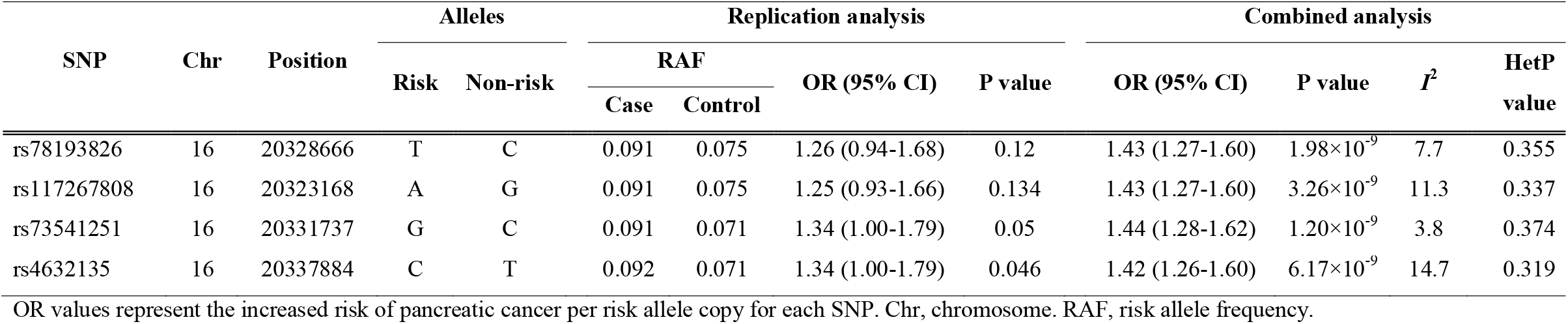
Associations between selected SNPs at 16p12.3 and pancreatic cancer risk in the replication and combined analyses

| SNP | Chr | Position | Alleles |  | Replication analysis |  |  |  | Combined analysis |  |  |  |
| --- | --- | --- | --- | --- | --- | --- | --- | --- | --- | --- | --- | --- |
| | | | Risk | Non-risk | RAF | | OR (95% CI) | P value | OR (95% CI) | P value | $I^2$ | HetP value |
|  |  |  |  |  | Case | Control |  |  |  |  |  |  |
| rs78193826 | 16 | 20328666 | T | C | 0.091 | 0.075 | 1.26 (0.94-1.68) | 0.12 | 1.43 (1.27-1.60) | $1.98 \times 10^{-9}$ | 7.7 | 0.355 |
| rs117267808 | 16 | 20323168 | A | G | 0.091 | 0.075 | 1.25 (0.93-1.66) | 0.134 | 1.43 (1.27-1.60) | $3.26 \times 10^{-9}$ | 11.3 | 0.337 |
| rs73541251 | 16 | 20331737 | G | C | 0.091 | 0.071 | 1.34 (1.00-1.79) | 0.05 | 1.44 (1.28-1.62) | $1.20 \times 10^{-9}$ | 3.8 | 0.374 |
| rs4632135 | 16 | 20337884 | C | T | 0.092 | 0.071 | 1.34 (1.00-1.79) | 0.046 | 1.42 (1.26-1.60) | $6.17 \times 10^{-9}$ | 14.7 | 0.319 |

OR values represent the increased risk of pancreatic cancer per risk allele copy for each SNP. Chr, chromosome. RAF, risk allele frequency.

We also examined the previously published pancreatic cancer risk loci from PanScan^14^, noting that 11 of those 19 SNPs were nominally statistically significant (rs13303010 at 1p36.33, rs3790844 at 1q32.1, rs9854771 at 3q28, rs2736098 at 5p15.33, rs401681 at 5p15.33, rs10094872 at 8q24.21, rs505922 at 9q34.2, rs9581943 at 13q12.2, rs7214041 at 17q24.3, and rs16986825 at 22q12.1) or genome-wide significant (rs9543325 at 13q22.1) in our GWAS meta-analysis (**Supplementary Table 6**). Notably, we confirmed the nominally significant association (*P*<3.84×10^-5^) between rs505922 of the ABO locus and pancreatic cancer risk.

Epidemiological studies have consistently shown that longstanding T2D is associated with an increased risk of pancreatic cancer^15^. Recently, 88 genetic variants, including both novel and established variants, were reported in a GWAS meta-analysis of T2D in the Japanese population^16^. We found that the top 3 SNPs at 16p12.3 (rs78193826, rs117267808, and rs73541251) were also genome-wide significantly associated with the risk of T2D in the latest GWAS comprising 191,764 Japanese subjects (**Supplementary Table 7**). In addition, these 3 SNPs were nominally significantly associated with hemoglobin A1c (HbA1c) (*P*<1×10^-4^) and blood glucose levels (*P*<0.01), which were included in another GWAS of quantitative traits in 42,790 and 93,146 Japanese subjects, respectively^17^. In addition to the *GP2* SNPs, 4 of the 82 T2D-associated SNPs (rs838720 at 2q37.1 (*DGKD*), rs2233580 at 7q32.1 (*PAX4*), rs2290203 at 15q26.1 (*PRC1-AS1*), and rs663129 at 18q21.32 (*MC4R*)) were nominally significant in our GWAS meta-analysis (**Supplementary Table 8**). In addition, one of the 25 HbA1c-associated SNPs (rs4728092 at 7q32.1 (*SND1*)) was nominally significant in our GWAS meta-analysis (**Supplementary Table 9**).

To examine whether the associations of T2D and T2D-related quantitative traits with pancreatic cancer are consistent with a causal effect, we performed a Mendelian randomization (MR) analysis with the inverse variance-weighted (IVW) and MR-Egger methods. No significant associations were observed between SNP-modulated T2D and pancreatic cancer based on the IVW method (**Figure 3a**). Instead, genetically increased HbA1c levels appeared to be causally related to an increased risk of pancreatic cancer, on the basis of the significant results with both the IVW and MR-Egger methods (**Figure 3b and Supplementary Figure 4b**).

**Figure 3.**
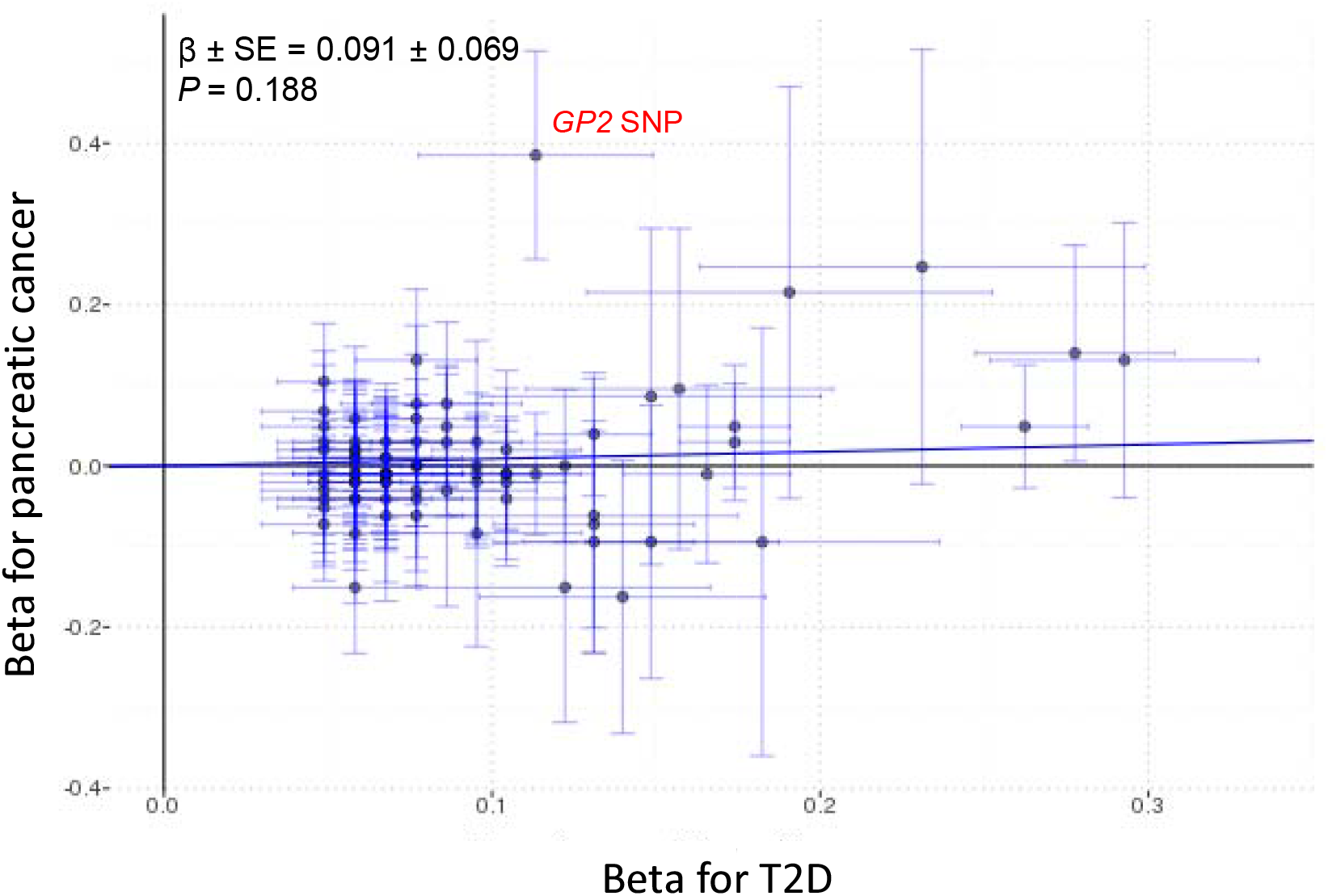
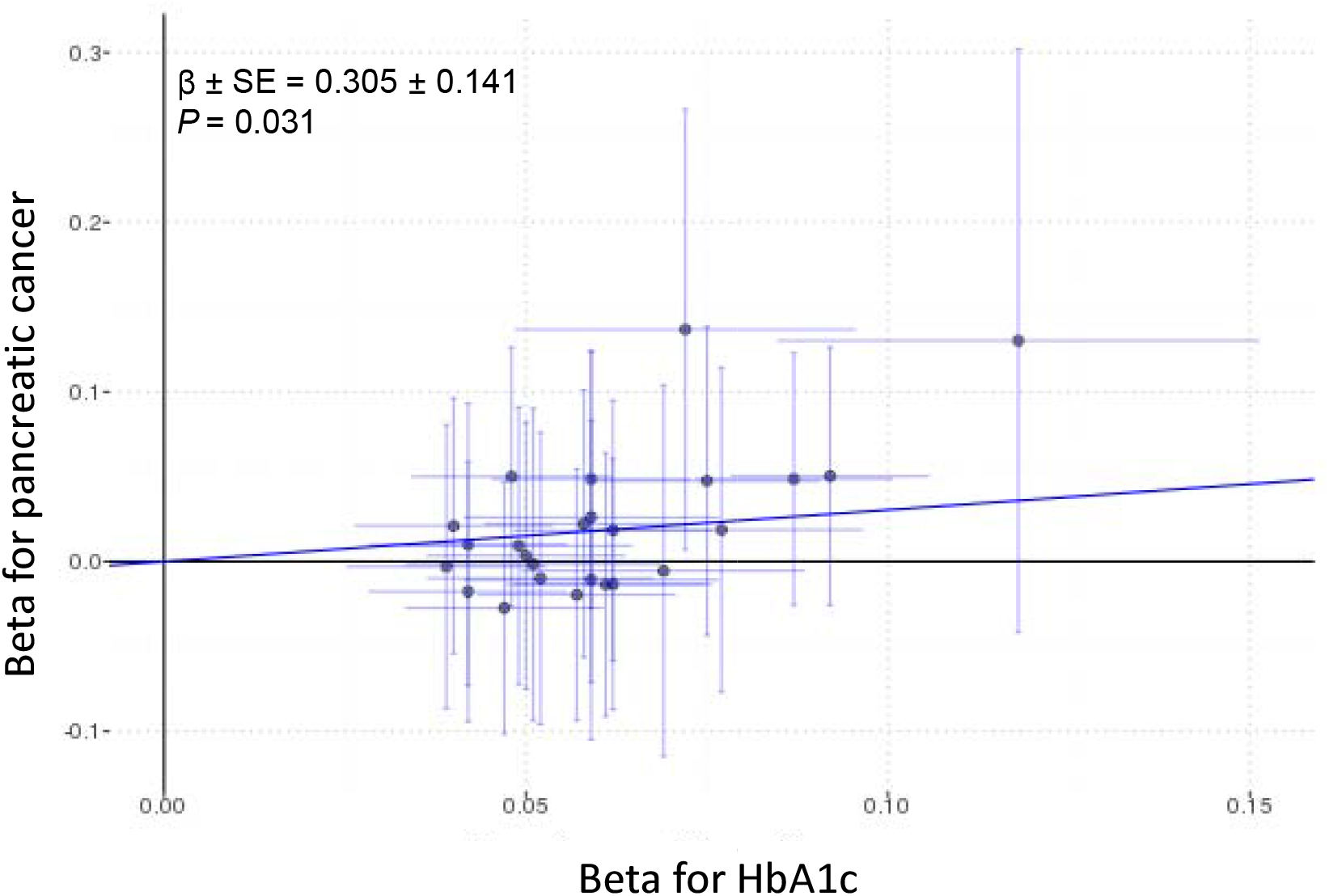
MR analysis with the IVW method for the relationship between T2D and pancreatic cancer in the Japanese population. (a) The results from 82 T2D-associated SNPs (b) 25 HbA1c-associated SNPs.

To complement the SNP-based GWAS, we performed a gene-based GWAS using MAGMA^18^ (**Supplementary Figure 5**). We confirmed the significant associations for *GP2* and *WNT2B* identified by the SNP-based GWAS. Notably, a novel significant association (Bonferroni-corrected *P* < 2.84 × 10^−6^) for the gene *KRT8* was observed (**Supplementary Table 10** and **Supplementary Figures 5 and 6**), and this association was further replicated in the PanScan 1 and PanScan 2 datasets (*P*=0.024)^13^.

## Discussion

The role of inherited common genetic variations in pancreatic cancer susceptibility remains incompletely understood. We identified and replicated a novel risk locus at 16p12.3 for pancreatic cancer through combining three GWAS datasets in the Japanese population. Furthermore, we provided evidence that the identification of this new locus can be attributed to the observed differences in the MAF of the lead SNP (rs78193826) at 16p12.3 and the LD structure in this region across ethnic populations.

Little overlap has been observed when risk loci reported from previous Chinese or Japanese GWASs are compared with those reported in the PanScan GWASs^2^. By including more than twice the number of cases than were included in previous Japanese or Chinese GWASs as well as imputed SNP data, we replicated the majority of the significant risk loci discovered in the PanScan GWASs (**Supplementary Table 6**). Moreover, for most variants, the direction and magnitude of the effects in our GWAS meta-analysis of Japanese subjects were consistent with those in populations of European ancestry. These findings suggested that GWAS-identified causal variants at many loci are shared across ancestral groups and that the lack of replication may be due to an insufficient sample size in previous Chinese or Japanese GWASs.

Several lines of evidence indicate that rs78193826 is most likely a causal variant at 16p12.3, which harbors the *GP2* gene. First, this variant is nonsynonymous; the nucleotide mutation from C to T causes an amino acid change from valine to methionine, which could affect protein structure and function. Second, functional annotations in several databases consistently indicate that this variant is highly pathogenic. Third, the observed differences in the MAF of rs78193826 as well as the LD structure across different ethnic populations provide indirect evidence supporting its role as a causal variant in the Japanese population. The frequency of the minor T allele of rs78193826 is 0.1% in populations of European ancestry but 7% in the Japanese population. Given this apparent difference in the MAF, rs78193826 could not have been identified in the PanScan GWASs, although the PanScan GWASs included a much larger sample size than our GWAS. Of the 10 SNPs in this region, only rs4383153 has association summary statistics available in PanScan publications; however, no significant associations were observed between this SNP and pancreatic cancer risk (**Supplementary Table 5**). While complete LD between rs4383153 and rs78193826 was evident in the Japanese population, no LD data were available for these two SNPs in the European ancestry populations (1000 Genomes Project Phase 3 CEU). As the frequency of the minor T allele of rs78193826 ranges from 3.9% to 6.6% in other Asian populations, rs78193826 is likely to be a causal variant for pancreatic cancer that is specific to Asian populations. However, further trans ethnicity replicability and fine mapping are necessary to establish the causal role of this variant.

Genetic variations in the *GP2* gene have been linked to several phenotypes in addition to pancreatic cancer. The SNP rs12597579, located in the upstream region of the *GP2* gene, has been associated with body mass index (BMI) in a GWAS including East Asians^19^. However, rs12597579 was not in LD with rs78193826 (r^2^=0.003, calculated from Japanese samples in the 1000 Genomes Project Phase 3), suggesting that rs12597579 may have functions different from those of rs78193826. Coincidently, the lead variant (rs117267808) in the *GP2* gene identified in the latest GWAS meta-analysis of T2D in the Japanese population is the same variant that we identified in our GWAS meta-analysis of pancreatic cancer (**Supplementary Table 7**). Of the 82 T2D-related SNPs, 5 showed significant associations (*P*<0.05) with pancreatic cancer, suggesting that pancreatic cancer and T2D may share specific genetic susceptibility factors. Furthermore, the risk alleles of rs78193826 and rs117267808 were identical for pancreatic cancer and T2D. Together, these findings indicate that *GP2* variants may exert pleiotropic effects on multiple traits.

The newly identified lead SNP (rs78193826) encodes the GP2 protein, which is present on the inner surface of zymogen granules in pancreatic acinar cells^20^. GP2 is a glycosylated protein of ~90 kDa that contains multiple sites, such as an asparagine-linked glycosylation site, a zona pellucida (ZP) domain, and a glycosylphosphatidylinositol (GPI) linkage to the membrane^21^. During the secretory process, GP2 is cleaved from the membrane and secreted into the pancreatic duct along with other digestive enzymes. The expression of GP2 is extremely high in normal pancreatic tissues compared to that in other tissues (**Supplementary Figure 7**)^22^. However, according to the GEPIA database, pancreatic tumor tissues have a decreased level of GP2 expression compared with that in nontumor tissues (**Supplementary Figure 8**)^23^. The functional consequence of the lead SNP can be, in part, inferred from its location in the *GP2* gene: the ZP domain. In addition to GP2, an increasing number of proteins, such as uromodulin^24^ and the transforming growth factor (TGF)-β receptor III^25^, have been found to contain a ZP domain, the functions of which involve extracellular matrix formation and polymerization^26^. In particular, uromodulin, the most abundant protein secreted by the kidney, shares 55% amino acid sequence identity with GP2 in the ZP-C subdomain^24^. Pathogenic mutations in the *UMOD* gene have been shown to affect the structure and polymerization of uromodulin, resulting in kidney disease^24^. Given the similarity in the amino acid sequence as well as the expected functionality between these two proteins, genetic variations in the ZP domain of *GP2* might also be associated with disease risk. Specifically, our GWAS finding that genetic variations in the *GP2* gene are associated with pancreatic cancer risk elucidates a possible role of bacterial infection in the pathogenesis of pancreatic cancer. The biological function of GP2 in the pancreas remains unclear; previous experiments did not show changes in exocrine secretion and morphology in *GP2* knockout mice^27^. However, accumulating evidence has demonstrated the effects of GP2 on the innate immune response^28,29^. The GP2 is also expressed in the membranous (M) cells of the intestinal epithelium in humans and mice, where it acts as an uptake receptor for a subset of commensal and pathogenic bacteria^30^. For example, GP2 and its closest homolog, uromodulin, have been shown to bind to *Escherichia coli (E. coli*) that express type 1 fimbriae^31^. In particular, uromodulin null mice showed increased sensitivity to urinary tract infections^24^. These findings suggest that GP2 may also play a role in host defense in the pancreas, given that proteobacteria have been detected in pancreatic ductal adenocarcinoma samples as well as in the normal human pancreas^32^.

Previous epidemiological studies have suggested that HbA1c levels, even in nondiabetic ranges, or changes in HbA1c levels in new-onset T2D are associated with pancreatic cancer risk^33, 34^ Our MR analysis provided corroborating evidence that genetically increased HbA1c levels may be causally associated with pancreatic cancer risk. This result was also partially consistent with a previous MR analysis, in which T2D was not causally implicated but BMI and fasting insulin were causally associated with pancreatic cancer^35^. Because known HbA1c-related genetic variants explain little of the variance in HbA1c levels, further studies are needed to strengthen the causal inference by incorporating more variants. The null findings on the causal role for T2D may reflect the phenotypic and genetic heterogeneity of T2D, but T2D may also be both a cause and consequence of pancreatic cancer^35^.

Three genome-wide significant genes (*GP2, WNT2B*, and *KRT8*) emerged in the gene-based GWAS. Among these genes, *WNT2B* was reported in the previous PanScan GWAS as a novel gene at 1p13.1 with suggestive evidence of association^6^, and *KRT8* is a novel finding. KRT8 belongs to a group of intermediate-filament cytoskeletal proteins involved in maintaining epithelial structural integrity^36^. KRT8 is expressed in both ductal and acinar single-layer epithelia, and mutations in the *KRT8* gene have been linked to exocrine pancreatic disorders and liver disease^37, 38^.

In conclusion, our GWAS meta-analysis identified a novel risk locus at chromosome 16p12.3, which harbors the *GP2* gene, for pancreatic cancer in the Japanese population. Further fine mapping and functional characterization are required to elucidate the effects of common *GP2* gene variants on pancreatic cancer susceptibility. Moreover, our findings highlight genetic susceptibility factors shared between T2D and pancreatic cancer.

## Methods

### Study samples

We performed a GWAS meta-analysis based on three Japanese studies: the Japan Pancreatic Cancer Research (JaPAN) consortium GWAS, the National Cancer Center (NCC) GWAS, and the BioBank Japan (BBJ) GWAS. An overview of the characteristics of the study populations is provided in **Supplementary Table 1.** Information on the study-specific genotyping, imputation, and analysis tools is provided in **Supplementary Table 2.** All studies were imputed based on the 1000 Genomes Project reference panel (Phase 3).

### JaPAN consortium GWAS

Participants in this GWAS were drawn from the JaPAN consortium^39^. Two case-control datasets were combined, resulting in a total of 945 pancreatic cancer cases and 3134 controls. The vast majority of cases were diagnosed as primary adenocarcinoma of the exocrine pancreas (ICD-O-3 codes C250–C259). The first dataset included 622 pancreatic cancer patients who were recruited from January 2010 to July 2014 at five participating hospitals in the Central Japan, Kanto, and Hokkaido regions. This multi-institutional case-control study collected questionnaire data on demographic and lifestyle factors, as well as 7-ml blood samples, from the study participants. The second dataset included 323 patients with newly diagnosed pancreatic cancer and 3134 control subjects recruited to an epidemiological research program at Aichi Cancer Center (HERPACC) between 2005 and 2012. All new outpatients on their first visit to Aichi Cancer Center were invited to participate in HERPACC. Those who agreed to participate filled out a self-administered questionnaire and provided a 7-ml blood sample. After quality control, 943 cases and 3057 controls remained for the subsequent analysis (**Supplementary Table 2**). None of the control subjects had a diagnosis of cancer at the time of recruitment. Written informed consent was obtained from all study participants, and the study protocol was approved by the Ethical Review Board of Aichi Medical University, the Institutional Ethics Committee of Aichi Cancer Center, the Human Genome and Gene Analysis Research Ethics Committee of Nagoya University, and the ethics committees of all participating hospitals.

### BBJ GWAS

Pancreatic cancer cases were obtained from the BBJ GWAS, which was launched in 2003 and collected DNA and clinical information from approximately 200,000 patients, including those with pancreatic cancer^40^. Overall, 422 pancreatic cancer cases with available genotype data were recruited from 2003 to 2008. Clinical information was collected using a standardized questionnaire. This study was approved by the ethics committees of the RIKEN Center for Integrative Medical Sciences. The controls were drawn from the participants in four population-based cohort studies in Japan: the Japan Multi-Institutional Collaborative Cohort Study (J-MICC), the Japan Public Health Center-based Prospective Study (JPHC), the Tohoku Medical Megabank Project Organization (ToMMo), and the Iwate Tohoku Medical Megabank Organization (IMM). A total of 28,870 controls who passed genotype data quality control assessments were included in the study. In all participating cohort studies, informed consent was obtained from the participants by following the protocols approved by the corresponding institutional ethics committees. The detailed descriptions of the BBJ and each cohort study are provided in **the Supplementary Note.**

### NCC GWAS

The case and control samples were derived from a previous pancreatic cancer GWAS^11^. The cases were 677 patients diagnosed with invasive pancreatic ductal adenocarcinoma at the NCC Hospital, Tokyo, Japan. The control population consisted of 677 Japanese volunteers who participated in a health checkup program in Tokyo. After preimputation quality control, 674 cases and 674 controls remained for the subsequent analysis (**Supplementary Table 2**). This project was approved by the ethics committee of the NCC.

### Quality control after genotype imputation

After genotype imputation, quality control was applied to each study. SNPs with an imputation quality of *r*^2^ < 0.5 or a MAF of <0.01 were excluded. SNPs that passed quality control in at least two cohorts were included in the meta-analysis.

### Association analysis for SNPs and pancreatic cancer

The association of pancreatic cancer with SNP allele dose was tested using logistic regression analysis with adjustment for the top 2 principal components. Other known covariates, such as age, sex, and cigarette smoking, were not included in the analysis because the inclusion of covariates has been shown to substantially reduce the power for the identification of disease-associated variants when the disease prevalence is less than 2%^41^.

The effect sizes and standard errors were used in the subsequent meta-analysis.

### Meta-analysis

We performed a meta-analysis of three pancreatic cancer GWASs (JaPAN, BBJ and NCC). The association results for each SNP across the studies were combined with METAL software in a fixed effects inverse variance-weighted meta-analysis. Heterogeneity in allelic effects was assessed using the *I*^2^ index. The meta-analysis included 7,914,378 SNPs with genotype data available from at least two cohorts. A *P* value threshold of 5 × 10^-8^ was used to determine genome-wide significance. We assessed the inflation of test statistics using the genomic control lambda.

### Replication analysis

The replication cohort comprised 507 cases and 879 controls who were recruited under the same framework as the multi-institutional case-control study of the JaPAN consortium (**Supplementary Table 1**). The cases and controls were recruited from August 2014 to March 2018 at five participating hospitals in the Central Japan, Kanto, and Hokkaido regions. The control subjects included outpatients as well as screening participants with no diagnosis of cancer. SNP genotyping was performed using a Fluidigm SNP assay at the Aichi Cancer Center Research Institute, with the laboratory staff blinded to the case-control status. The association of pancreatic cancer with SNP allele dose was tested using unadjusted logistic regression analysis.

### Functional annotations

To prioritize the associated SNPs at the novel loci, we adopted a series of bioinformatic approaches to collate functional annotations. We first used ANNOVAR^42^ to obtain an aggregate set of functional annotations—including the gene location and the impact of the amino acid substitution based on prediction tools such as SIFT, PolyPhen-2, and CADD—for SNPs with a *P* value of <5 × 10^−8^ for pancreatic cancer. We also explored potential effects on gene regulation by annotating these SNPs using the RegulomeDB database^43^.

### MR analysis

We performed MR analyses using independent, genome-wide significant T2D-associated or HbA1c-associated SNPs, which were available from two published GWAS meta-analyses in Japanese subjects, as instrumental variables^16,17^. For the two-sample MR analysis of T2D and pancreatic cancer, we did not exclude the overlapping samples (15.5% found only in the controls) because retaining these samples was unlikely to introduce substantial bias^44^. A total of 106 pancreatic cancer cases were excluded in the HbA1c GWAS, and the effect sizes for the HbA1c-associated SNPs were reestimated. After the exclusion of 6 SNPs on the X chromosome (5 SNPs for T2D and 1 SNP for HbA1c) and an SNP for T2D (rs35678078) without genotype data, the summary data for 82 T2D-related SNPs and 25 HbA1c-related SNPs and the associations of these SNPs with pancreatic cancer risk were analyzed using IVW and MR-Egger regression methods. MR analysis was performed with the MendelianRandomization package^45^.

### Gene-based analysis

SNP-based *P* values were combined into gene-based *P* values using MAGMA software version 1.06^18^. SNP summary statistics (*P* values) from the meta-analysis were used as input for MAGMA. In gene-based association tests, LD between SNPs was accounted for, and the *P* value threshold for genome-wide significant associations was set at 2.84×10^−6^. The 1000 Genomes reference panel (Phase 3, East Asian) was used to control for LD. We did not include any upstream/downstream regions around the genes in this analysis; only variants located between the first exon and the last exon of a gene were used to calculate the gene-based *P* values. The NCBI Gene database was used to define genomic intervals for protein-coding genes. To replicate the association between *KRT8* and pancreatic cancer, we applied SNP summary statistics from PanScan 1 and PanScan 2 (pha002889.1)^13^ to MAGMA. The MAF of the Haplotype Map (HapMap) project Phase 2 CEU samples for each SNP was added to the summary statistics because the pha002889.1 data did not include the MAFs. We excluded variants with a call rate (CR) of > 95% in the cases, a CR of > 95% in the controls, a Hardy-Weinberg equilibrium test *P* value of <1 × 10^−6^ in the controls, and/or a MAF of <0.01. The 1000 Genomes reference panel (Phase 3, European) was used to control for LD. The significance level was set at α=0.05.

## Acknowledgments

We thank Mayuko Masuda, Kikuko Kaji, Kazue Ando, Etsuko Ohara, Sumiyo Asakura, and Keiko Hanai for assistance with data collection. We thank Yumiko Kasugai for assistance with SNP genotyping. We would such as to express our gratefulness to the staff of BioBank Japan for their outstanding assistance.

This work was supported by the Ministry of Health, Labor, and Welfare of Japan (H21-11-1) and the Ministry of Education, Culture, Sports, Science, and Technology of Japan (Nos. 16H06277, 17K09095, 26253041, and 17015018). HERPACC, a part of the JaPAN consortium, was supported by Grants-in-Aid for Scientific Research for Priority Areas of Cancer (No. 17015018) and Innovative Areas (No. 221S0001) and JSPS KAKENHI grants (Nos. 26253041, 15H02524, 16H06277, and 18H03045) from the Japanese Ministry of Education, Science, Sports, Culture and Technology; and a Grant-in-Aid for the Third Term Comprehensive 10-year Strategy for Cancer Control from the Ministry of Health, Labor and Welfare of Japan and the Cancer Biobank Aichi.

This study was partially supported by the BioBank Japan project and the Tohoku Medical Megabank project, which is supported by the Ministry of Education, Culture, Sports, Sciences and Technology of Japan and the Japan Agency for Medical Research and Development.

The JPHC Study has been supported by the National Cancer Center Research and Development Fund since 2011 and was supported by a Grant-in-Aid for Cancer Research from the Ministry of Health, Labor and Welfare of Japan from 1989 to 2010.

The J-MICC Study was supported by Grants-in-Aid for Scientific Research for Priority Areas of Cancer (No. 17015018) and Innovative Areas (No. 221S0001) and JSPS KAKENHI Grant (No. 16H06277) from the Japanese Ministry of Education, Science, Sports, Culture and Technology.

## Author contributions

Y.L., M.N., H.I., A.I., M.O., T.O., S.Kamiya., and C.W. designed the study. Y.L, M.N., and Y. Kamatani wrote the manuscript. M.N., F.K., Y. Kobayashi., Y.Kamatani, M.A. K.I. performed the statistical analysis. H.S., H.I., M.O., T.S., M.Matsuyama., N.S., M.Morimoto., S.Kobayashi., T.F., M.U., S.O., N.E., S.Kuruma. M.Mori., H.N., Y.A., K.H., Y.S., Y.M., K.M., M.Hirata., K.Shimada., T.O, Y.W., K.Kuriki., A.K., K.W., T.Yamaji., M.I., N.S., S.T., K.Kinoshita., N.F., F.K., A.S., S.N., K.T., K.Suzuki., Y.O., M.Horikoshi., T.Yamauchi., T.Kadowaki., T.Yoshida. contributed to the data acquisition. A.I., T.M., Y.Hayashi., Y.Hosono., H.E., T. Kohmoto., I.I., M.K. T.Kawaguchi., M.T., and F.M. contributed to the SNP genotyping. S.Kikuchi. and K.M. supervised the study. All authors approved the final version of the manuscript.

## Completing financial interests

The authors declare no competing financial interests.

## Data availability statement

Summary data will be available from a public data repository. Restrictions apply to the availability of these data; however, data are available from the authors upon reasonable request.

