## Supplementary information for "Genome-wide association meta-analysis identifies novel *GP2* gene risk variants for pancreatic cancer in the Japanese population"

Supplementary Tables

Supplementary Table 1. Characteristics of the study participants 　 p. 3

Supplementary Table 2. Cohort-specific information on genotyping, imputation

and association testing 　p. 4 (Electronic Excel file)

Supplementary Table 3. Genomic region and lead SNPs associated with

pancreatic cancer susceptibility in each of three Japanese GWASs

　p. 4 (Electronic Excel file)

Supplementary Table 4. Allele frequency, effect size, and functional annotation for 10

SNPs with genome-wide significance at 16p12.3

p. 4 (Electronic Excel file)

Supplementary Table 5. Association results for rs4383153, located at 16p12.3, in the

PanScan GWAS 　　　　　　p. 5

Supplementary Table 6. Associations of pancreatic cancer with 19 SNPs with

genome-wide significance in the previous GWAS reported by Klein et al.

p. 6 (Electronic Excel file)

Supplementary Table 7. Associations between the top 3 SNPs for pancreatic cancer and T2D risk and quantitative traits in the Japanese GWAS

　　　　　　　　　　　　　　　　 p. 6 (Electronic Excel file)

Supplementary Table 8. The T2D SNPs most recently identified in the Japanese

population by a GWAS (Suzuki et al., Nat Genet, in press) and the associations

of these SNPs with pancreatic cancer risk in the present GWAS meta-analysis

p. 6 (Electronic Excel file)

Supplementary Table 9. The HbA1c SNPs most recently identified in the Japanese

population by a GWAS (Kanai et al., Nat Genet, 2017) and the associations

of these SNPs with pancreatic cancer risk in the present GWAS meta-analysis

p. 7 (Electronic Excel file)

Supplementary Table 10. Results of the gene-based tests using MAGMA

p. 7 (Electronic Excel file)

Supplementary Figures

Supplementary Fig. 1 Q-Q plot for the P values in the meta-analysis. p. 8

Supplementary Fig. 2 Regional association plots for the six loci identified in the meta-analysis. p. 9-15

Supplementary Fig. 3 LD maps of 10 SNPs with genome-wide significance at

16p12.3 based on 1000 Genomes (a) JPT and (b) CEU subjects. p. 16

Supplementary Fig. 4 MR analysis with the MR-Egger method of the relationship between T2D and pancreatic cancer in the Japanese population. p. 17-18

Supplementary Fig. 5 Manhattan plot for the gene-based analysis. p. 19

Supplementary Fig. 6 Q-Q plot for the P values in the gene-based analysis. p. 20

Supplementary Fig. 7 GP2 differential gene expression illustrated across tissues. p. 21

Supplementary Fig. 8 Comparison of the GP2 expression levels between normal and tumor samples.

p. 22

Supplementary Note

Additional details on the Biobank Japan Project and the population-based cohort studies p. 23-27

**Supplementary Table 1. Characteristics of the study participants**

| Phase | Study name | Group | Source | Sample size^a^ | Age (Mean±SD) | Male (%) |
| --- | --- | --- | --- | --- | --- | --- |
| GWAS | JaPAN | Case | Hospital | 943 | 64.7 ± 10.1 | 62.5 |
|  |  | Control | Hospital, Screening facility | 3,057 | 52.1 ± 11.8 | 49.3 |
|  | NCC | Case | Hospital | 674 | 62.7 ± 9.2 | 60.2 |
|  |  | Control | Volunteers, Health checkup program | 674 | 43.6 ± 10.0 | 63.6 |
|  | BBJ^b^ | Case | Hospital | 422 | 66.3 ± 10.0 | 66.4 |
|  |  | Control | Population-based cohort participants | 28,861 | 56.3 ± 10.0 | 39.4 |
| Replication | JaPAN | Case | Hospital | 507 | 66.3 ± 9.1 | 53.3 |
|  |  | Control | Hospital, Screening facility | 879 | 61.8 ± 11.2 | 51.2 |

^a^ Sample size indicates the number of samples that passed quality control and were subjected to genome-wide meta-analysis.

^b^ In the BBJ GWAS, control subjects were recruited from population-based cohort studies, including J-MICC, JPHC, ToMMo, and IMM.

**Supplementary Table 2. Cohort-specific information on genotyping, imputation and association testing**

#SNPs is the number of autosomal SNPs.

Enclosed electronic Excel file

**Supplementary Table 3. Genomic region and lead SNPs associated with pancreatic cancer susceptibility in each of three Japanese GWASs.**

*OR* values represent the increased risk of pancreatic cancer per risk allele copy for each SNP. Chr, chromosome. RAF, risk allele frequency.

Enclosed electronic Excel file

**Supplementary Table 4. Allele frequency, effect size, and functional annotation for 10 SNPs with genome-wide significance at 16p12.3**

*OR* values represent the increased risk of pancreatic cancer per risk allele copy for each SNP. Chr, chromosome. RAF, risk allele frequency.

Enclosed electronic Excel file

**Supplementary Table 5. Association results for rs4383153, located at 16p12.3, in the PanScan GWAS**

| **SNP** | **Chr** | **Position** | **Alleles** | | **Study (dbGap accession)** | **Reference** | **N** | | ***OR* (95% CI)** | **P value** |
| --- | --- | --- | --- | --- | --- | --- | --- | --- | --- | --- |
|  |  |  | **Risk** | **Non-risk** |  |  | **Case** | **Control** |  |  |
| rs4383153 | 16 | 20338622 | A | G | PanScan 1 (pha002874.1) | Amundadottir et al. (Nat Genet 41:986 (2009)) | 1896 | 1939 | 0.99 (0.71-1.39) | 0.962 |
|  |  |  |  |  | PanScan 1 and PanScan 2 (pha002889.1) | Li et al. (Carcinogenesis 33:1384 (2012)) | 3851 | 3934 | 1.04 (0.83-1.32) | 0.725 |

*OR* values represent the increased risk of pancreatic cancer per risk allele copy for each SNP. Chr, chromosome.

**Supplementary Table 6. Associations of pancreatic cancer with 19 SNPs with genome-wide significance in the previous GWAS reported by Klein et al.**

Although Klein et al. reported 22 SNPs in Table 1 and Supplementary Table 2 of their paper, rs35226131, rs73328514, and rs7190458 were not included in our meta-analysis.

Risk alleles increase the risk of pancreatic cancer in the Klein et al. study.

Meta-analysis results were not available for rs13303010 in NCC or for rs7214041 in JaPAN.

*OR* values represent the increased risk of pancreatic cancer per risk allele copy for each SNP. Chr, chromosome. RAF, risk allele frequency.

Enclosed electronic Excel file

**Supplementary Table 7. Associations between the top 3 SNPs for pancreatic cancer and T2D risk and quantitative traits in the Japanese GWAS**

*OR* values represent the increased risk of T2D per risk allele copy for each SNP. Chr, chromosome. RAF, risk allele frequency.

Beta values represent the change in the rank-based inverse normal transformed values of HbA1c and the blood glucose levels per risk allele copy for each SNP.

Enclosed electronic Excel file

**Supplementary Table 8. The T2D SNPs most recently identified in the Japanese population by a GWAS (Suzuki et al., Nat Genet, in press) and the associations of these SNPs with pancreatic cancer risk in the present GWAS meta-analysis**

Risk alleles increase the risk of T2D in the Japanese GWAS.

*OR* values represent the increased risk of T2D or pancreatic cancer per risk allele copy for each SNP. Chr, chromosome. RAF, risk allele frequency.

Meta-analysis results were not available for rs2233580 and rs28624681 in NCC or for rs12454712 in JaPAN.

Enclosed electronic Excel file

**Supplementary Table 9. The HbA1c SNPs most recently identified in the Japanese population by a GWAS (Kanai et al., Nat Genet, 2017) and the associations of these SNPs with pancreatic cancer risk in the present GWAS meta-analysis**

Risk alleles increase the HbA1c level in the Japanese GWAS.

*OR* values represent the increased risk of pancreatic cancer per risk allele copy for each SNP.

Beta values represent the change in the rank-based inverse normal transformed values of the HbA1c level per risk allele copy for each SNP. Chr, chromosome. RAF, risk allele frequency.

Enclosed electronic Excel file

**Supplementary Table 10. Results of the gene-based tests using MAGMA**

Genome-wide results are displayed in bold (P<2.84E-6 for the Bonferroni correction for the number of genes).

Enclosed electronic Excel file

**Supplementary Figure 1. Q-Q plot for the P values in the meta-analysis.** The vertical and horizontal axes indicate the observed and expected –log_10_(P value) for tests of association between SNPs and pancreatic cancer, respectively.

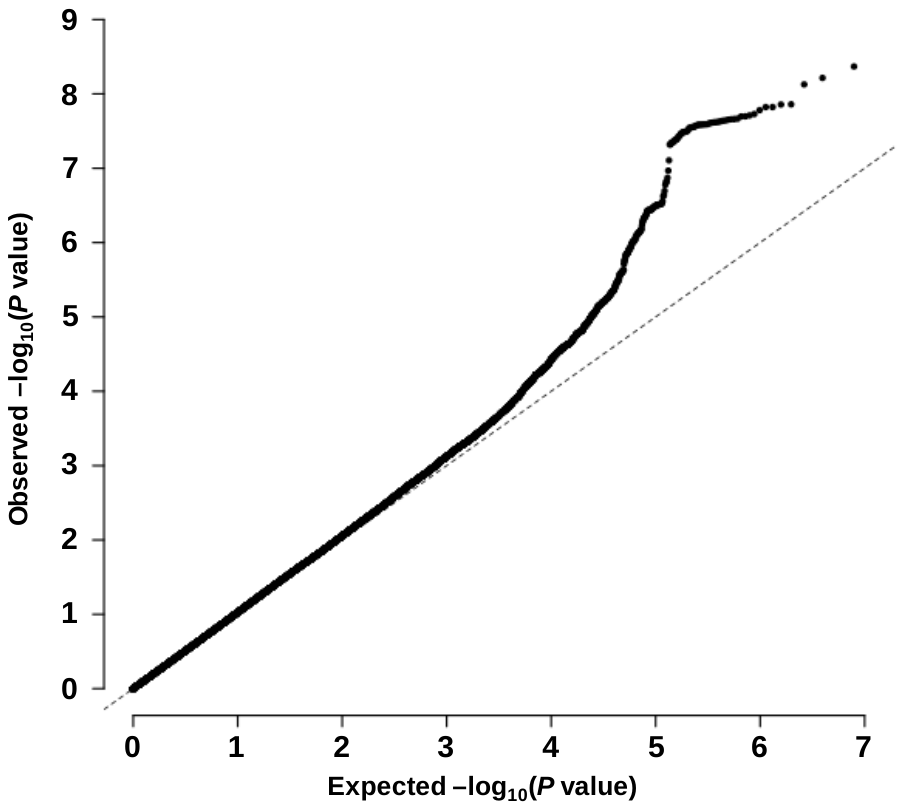

**Supplementary Figure 2. Regional association plots for the six loci identified in the meta-analysis.** The vertical axis indicates the –log_10_(P value) for the assessment of the association of each SNP with pancreatic cancer. Panels a to f show the plots for chromosome (chr) 1p13.2, 2p12, 3p12.3, 9q34.2, 13q12.2, or 13q22.1, in order. The colors indicate the LD (r^2^) between each sentinel SNP and neighboring SNPs based on the JPT population in the 1000 Genomes Project Phase 3.

**a**

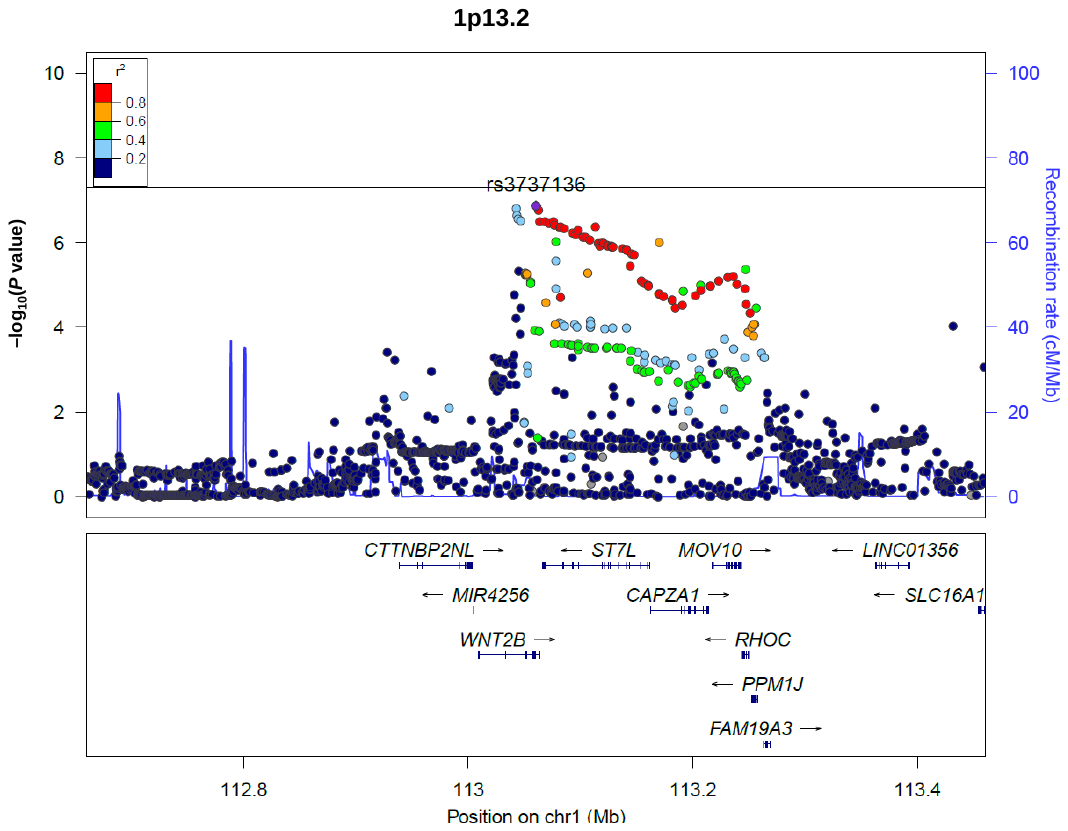

**b**

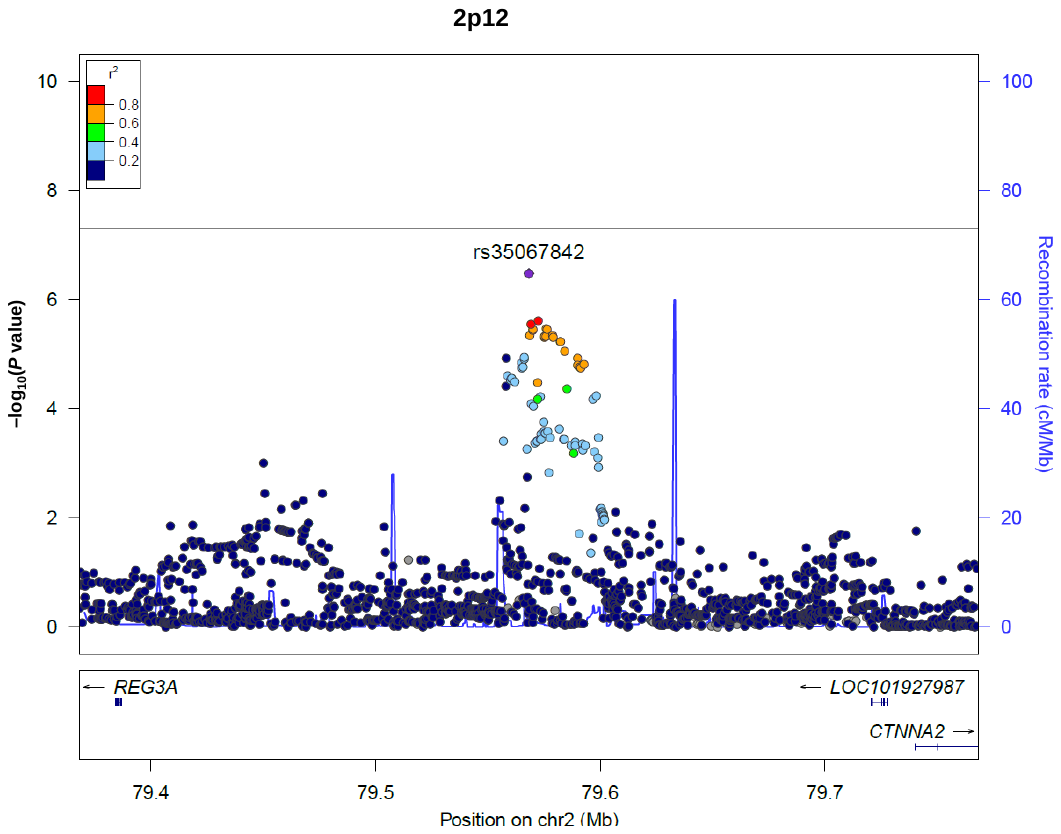

**c**

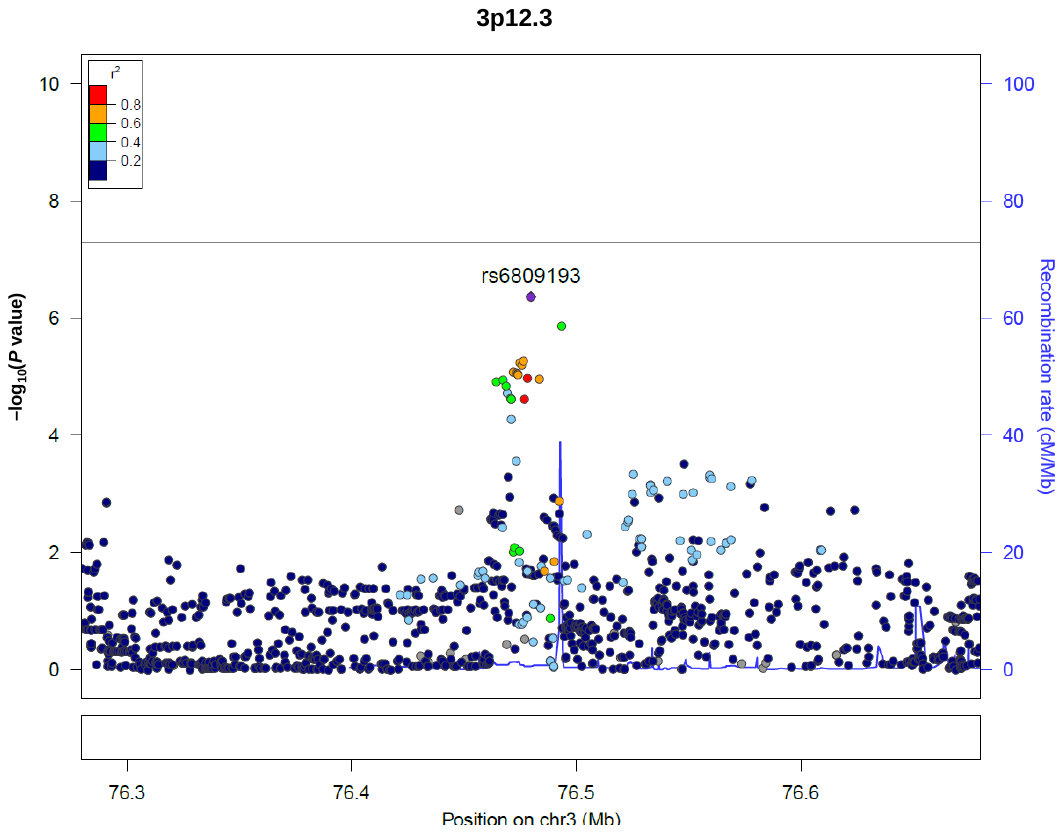

**d**

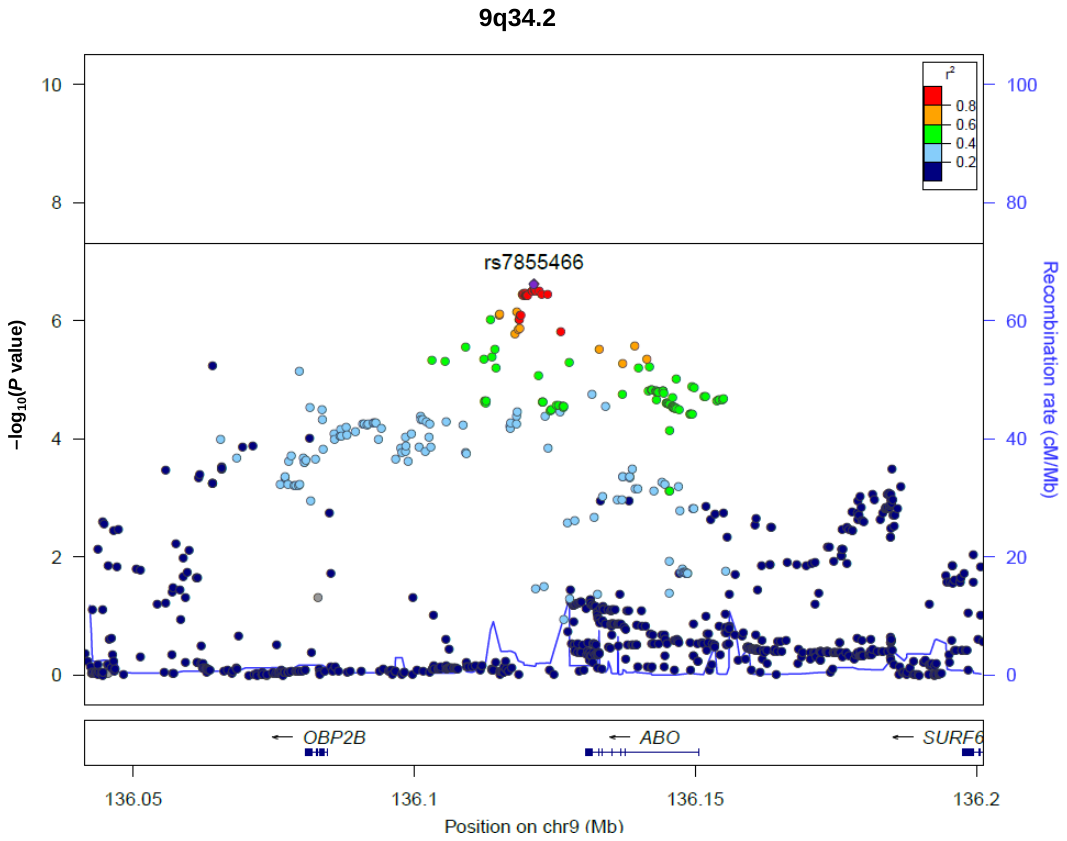

**e**

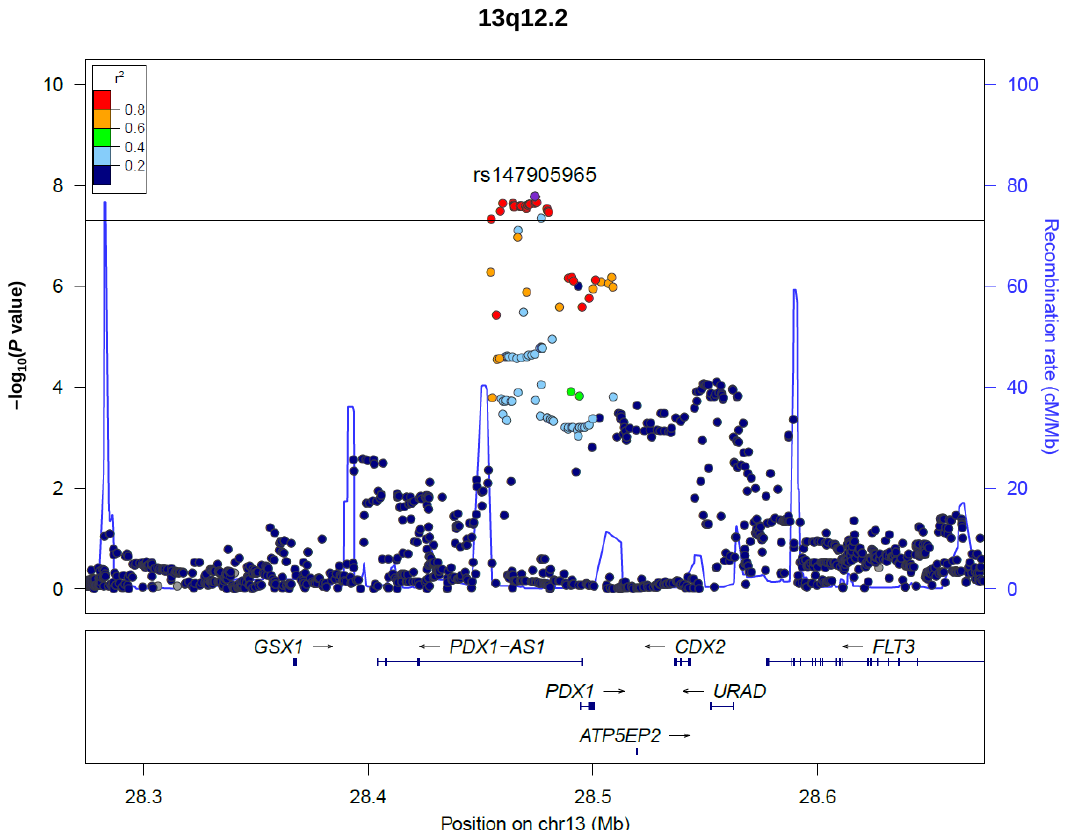

**f**

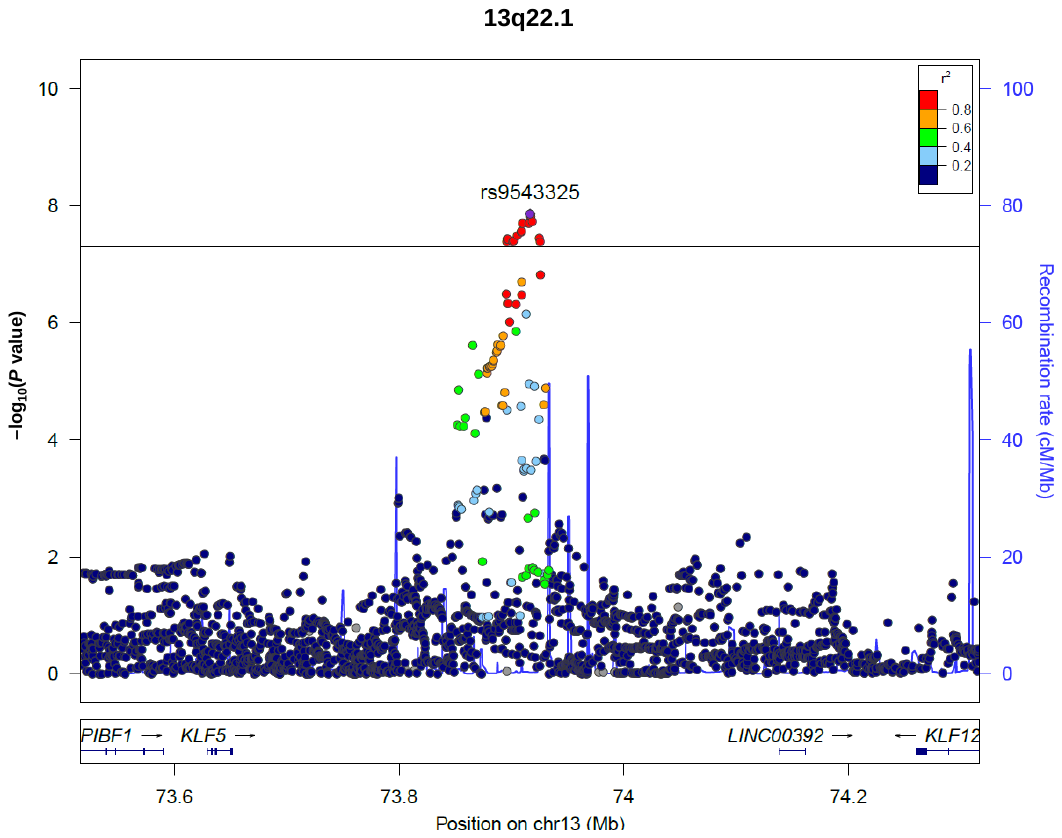

**Supplementary Figure 3. LD maps of 10 SNPs with genome-wide significance at 16p12.3 based on 1000 Genomes**

Pairwise linkage disequilibrium r^2^ values (white to black scales indicate low to high values), as determined with Haploview

(a) **JPT subjects　　　　　　　　　　　　　　　　　　　　　　　　　 (b) CEU subjects.**

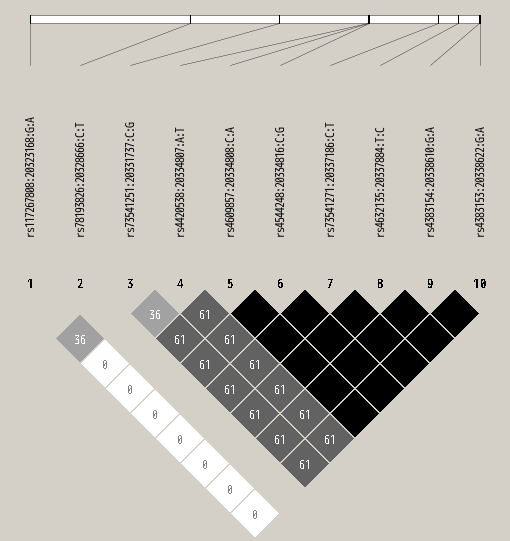

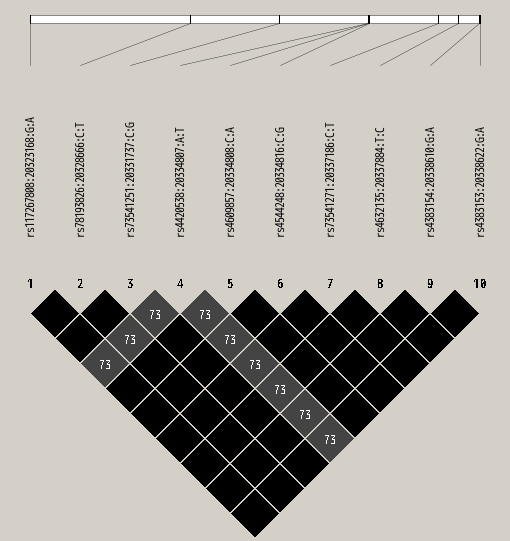

**Supplementary Figure 4.** MR analysis with the MR-Egger method of the relationship between T2D and pancreatic cancer in the Japanese population. (a) The results for 82 T2D-associated SNPs (b) the results for 25 HbA1c-associated SNPs.

**a**

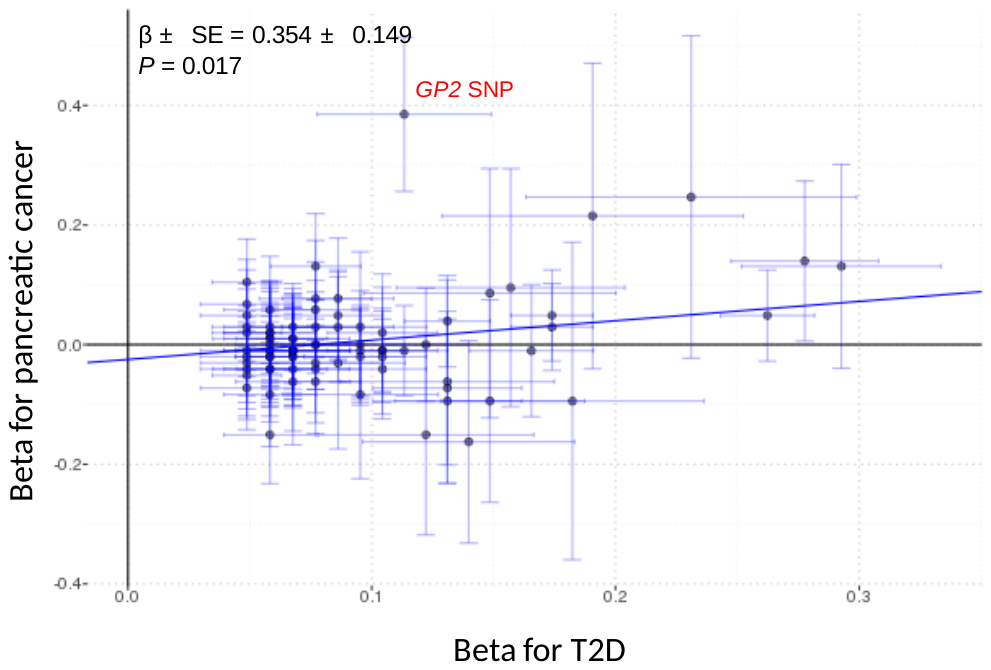

**b**

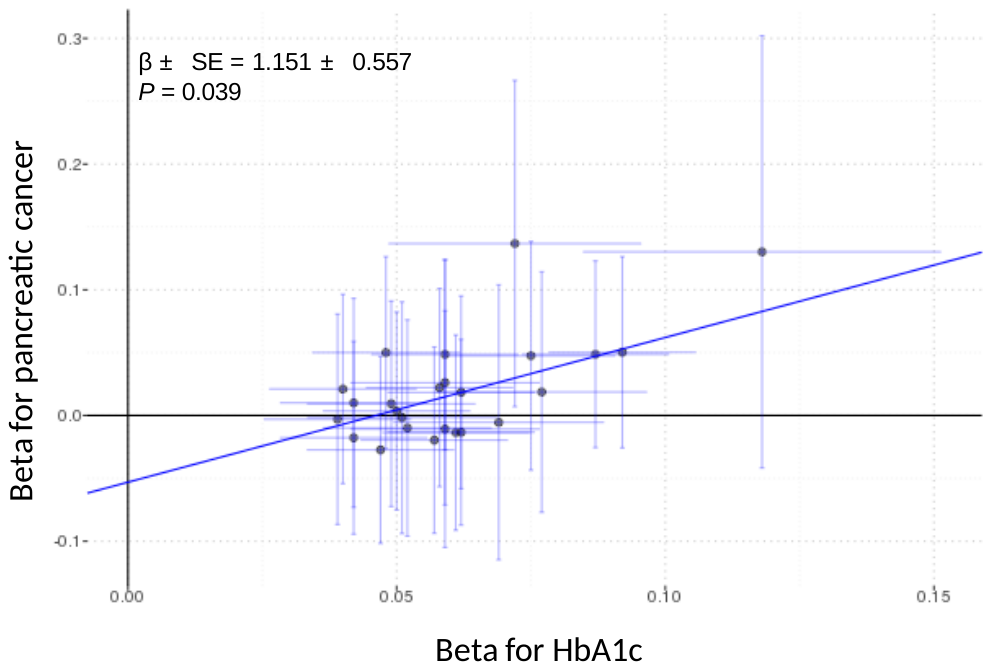

**Supplementary Figure 5. Manhattan plot for the gene-based analysis**. The horizontal red line represents the genome-wide significance level (α = 2.84 × 10^−6^).

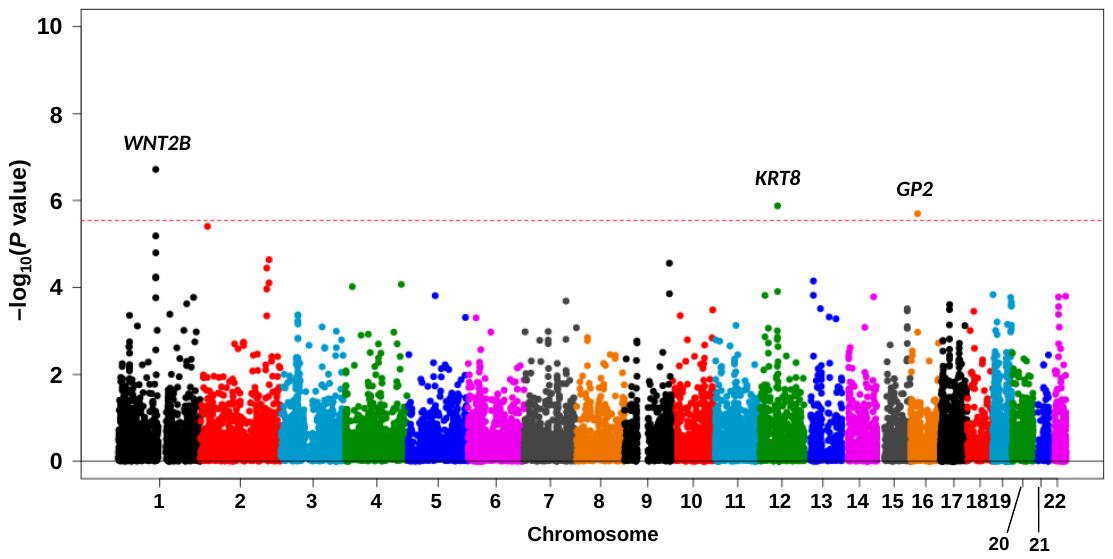

**Supplementary Figure 6. Q-Q plot for the *P* values in the gene-based analysis.** The vertical and horizontal axes indicate the observed and expected –log_10_(*P* value) for the tests of association between genes and pancreatic cancer.

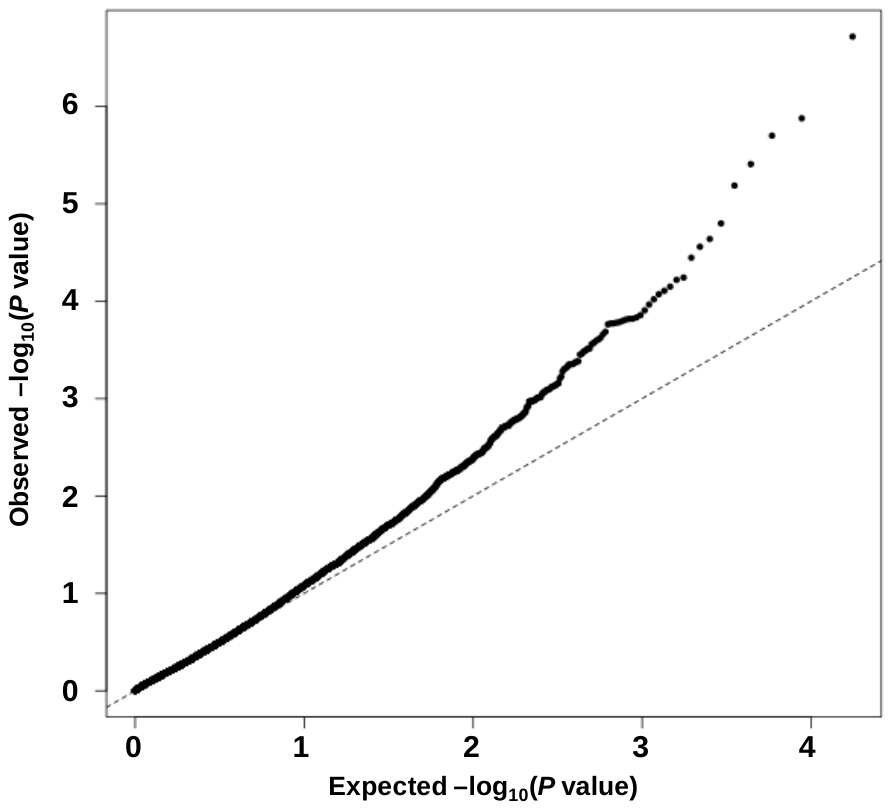

**Supplementary Figure 7. GP2 differential gene expression illustrated across tissues**. The vertical axis indicates the TPM (transcripts per million) value, and the horizontal axis shows the tissues. The data used for the analyses described in this manuscript were obtained from the GTEx portal on 09/16/18.

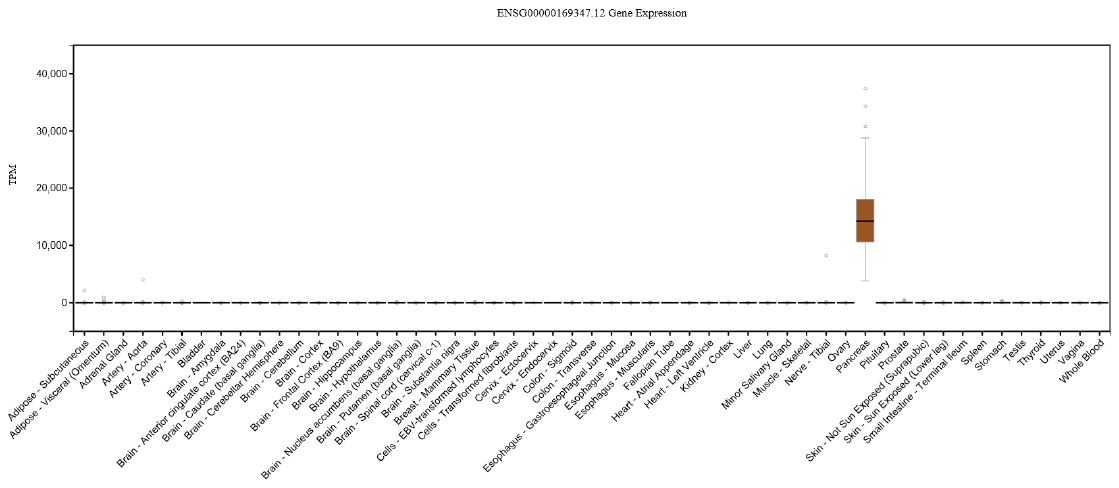

**Supplementary Figure 8. Comparison of GP2 expression levels between the normal and tumor samples.** The data were obtained from the GEPIA portal. The vertical axis indicates the log-transformed TPM (transcripts per million) value.

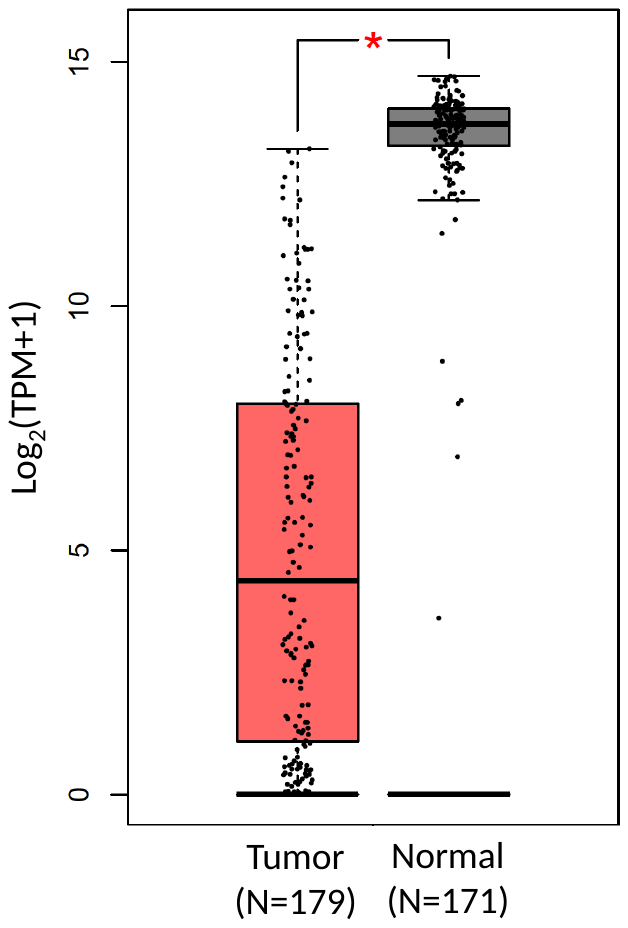

**Additional details on the BioBank Japan Project** The Biobank Japan (BBJ, http://biobankjp.org) project was started in 2003 and collected DNA and clinical information from a total of 200,000 patients with at least one of 47 common diseases, including pancreatic cancer. These subjects were recruited from a collaborative network of 66 hospitals organized by 12 medical institutions in Japan (Osaka Medical Center for Cancer and Cardiovascular Diseases, the Cancer Institute Hospital of the Japanese Foundation for Cancer Research, Juntendo University, Tokyo Metropolitan Geriatric Hospital, Nippon Medical School, Nihon University School of Medicine, Iwate Medical University, Tokushukai Hospitals, Shiga University of Medical Science, Fukujuji Hospital, National Hospital Organization Osaka National Hospital, and Iizuka Hospital). The eligibility of the patients was determined by physicians at the hospitals. Overall, 422 patients with pancreatic cancer for whom genotype data were available were recruited from BBJ for this study. Clinical information was collected via a standardized questionnaire through a medical records survey. For the controls, we used quality control-accepted genotype data for 28,861 individuals from four population-based studies: the Japan Multi-Institutional Collaborative Cohort Study (J-MICC), the Japan Public Health Center-based Prospective Study (JPHC), the Tohoku Medical Megabank Project Organization (ToMMo), and the Iwate Tohoku Medical Megabank Organization (IMM). A separate manuscript with the results of this single-association analysis, along with analyses of 43 additional diseases, is under review (Ishigaki et al.)

**Japan Multi-Institutional Collaborative Cohort Study (J-MICC study).** In the J-MICC study, 40,892 men and 51,750 women aged 35 to 69 years completed medical history questionnaires and donated blood samples at the time of the baseline survey, between 2004 and 2014^1^. The participants were recruited in 14 study areas throughout Japan among community dwellers, patients at the first visit to a cancer hospital, and health checkup examinees. For the present analyses, approximately 500 to 2,000 participants were selected from each study area, considering the number of respondents from each field and the geographical distribution of the subjects. All participants provided written informed consent. The ethics committees of Nagoya University (the affiliation of the principal investigator) and the other participating institutions approved the protocol for the J-MICC study. The following research institutions participated in the study: Chiba Cancer Center, University of Shizuoka, Nagoya City University, Aichi Cancer Center, Nagoya University, Shiga University of Medical Science, Tsuruga Nursing University, Kyoto Prefectural University of Medicine, University of Tokushima, Kyushu University, Saga University, and Kagoshima University.

**Japan Public Health Center-based Prospective Study (JPHC).** The JPHC samples were derived from a cohort of 33,736 residents in 9 public health center (PHC) areas who not only returned a self-administered questionnaire but also donated 10 mL of venous blood at the time of the baseline survey^2^. For the first sample selection step, we stratified the cohort by sex, 5-year age categories, and 9 PHC areas and then conducted random sampling, in which a similar proportion of subjects was selected from each stratum. Consequently, we determined 9,296 subjects for inclusion in the present GWAS. Before using the JPHC samples for genetic research, we obtained approval from the institutional review board of the National Cancer Center (Approval No.: 2011-044), Tokyo, Japan, and provided all eligible subjects the opportunity to refuse participation in the research.

**The Tohoku Medical Megabank (TMM) Project (Tohoku Medical Megabank Organization (ToMMo) and Iwate Tohoku Medical Megabank Organization (IMM).**

The TMM project is a reconstruction project from the Great East Japan Earthquake (2011) conducted by Tohoku University (http://www.megabank.tohoku.ac.jp/english/) and Iwate Medical University (http://iwate-megabank.org/en/)^3^. The TMM project encompasses two prospective cohort studies in Miyagi and Iwate Prefectures, Japan: the TMM Community-Based Cohort Study (TMM CommCohort Study) and the TMM Birth and Three-Generation Cohort Study (TMM BirThree Cohort Study). The TMM CommCohort Study is a population-based adult cohort study and recruited approximately 84,000 participants aged 20 years or older during 2013–2016. As of July 2017, the TMM BirThree Cohort Study had recruited approximately 74,000 participants, including fetuses and their parents, siblings, grandparents, and extended family members. All participants in the TMM project consented to genetic studies. Biospecimens (blood and urine) and medical data (questionnaires, blood and urine tests, and physiological measurements) were collected at the baseline examination. These samples and information are stored in the integrated biobank of the TMM project. DNA samples of the participants in the TMM CommCohort Study recruited in 2013 were analyzed by using the Illumina OmniExpressExome array (N=10,000). Information about age and sex was collected by using self-administered questionnaires and by reviewing municipal basic resident registers. Of the 10,000 persons with available genotype data, 9,202 had their height and weight measured in a standard manner. For persons without the body height and weight measurements (N=798), values of these variables were obtained from self-reported questionnaires when available (N=703). The remaining 95 persons who had neither measured nor self-reported values were excluded from the analyses.
